# Robotic selection for the rapid development of stable CHO cell lines for HIV vaccine for production

**DOI:** 10.1101/317537

**Authors:** Sara M O’Rourke, Gabriel Byrne, Gwen Tatsuno, Meredith Wright, Bin Yu, Kathryn A Mesa, Rachel C Doran, David Alexander, Phillip W Berman

**Affiliations:** Department of Biomolecular Engineering, The University of California at Santa Cruz, Santa Cruz, CA

**Keywords:** HIV vaccine, rgp120, carbohydrate, gene editing, CHO cells, antibodies, neutralization, robotic selection

## Abstract

The production of envelope glycoproteins (Envs) for use as HIV vaccines is challenging. The yield of Envs expressed in stable Chinese Hamster Ovary (CHO) cell lines is typically 10-100 fold lower than other glycoproteins of pharmaceutical interest. Moreover, Envs produced in CHO cells are typically enriched for sialic acid containing glycans compared to virus associated Envs that possess mainly high-mannose carbohydrates. This difference alters the net charge and biophysical properties of Envs and impacts their antigenic structure. Here we employ a novel gene-edited CHO cell line (MGAT1^-^ CHO) to address the problems of low expression, high sialic acid content, and poor antigenic structure. We demonstrate that stable cell lines expressing high levels of gp120, potentially suitable for biopharmaceutical production can be created using the MGAT1^-^ CHO cell line. We also show that the efficiency of this process can be greatly improved with robotic selection. Finally, we describe a MGAT1^-^ CHO cell line expressing A244-rgp120 that exhibits improved binding of three major families of bN-mAbs compared to Envs produced in normal CHO cells. The new strategy described has the potential to eliminate the bottleneck in HIV vaccine development that has limited the field for more than 25 years.

## 1 Introduction

The development of a safe, effective, and affordable HIV vaccine is a global public health priority. After more than 30 years of HIV research, a vaccine with these properties has yet to be described. To date, the only clinical study to show that vaccination can prevent HIV infection is the 16,000-person RV144 trial carried out in Thailand between 2003 and 2009 (1). This study involved immunization with a recombinant canarypox virus vector to induce cellular immunity (2-4) and a bivalent recombinant gp120 vaccine designed to elicit protective antibody responses (5-7). Although statistically significant, the protective efficacy of this vaccination regimen was low (31.2%, P=0.04). Several correlates of protection studies suggested that the protection observed was primarily due to antibodies to rgp120 (8-10). Thus, there is considerable interest in finding ways to improve the level of protection that can be achieved with rgp120 vaccine regimens. Improving an existing vaccine such as RV144, with an established record of safety, would be faster and more cost-effective than developing a new vaccine concept from scratch. A roadmap to improve the rgp120 vaccine used in the RV144 trial has been provided by the recent studies of broadly neutralizing monoclonal antibodies (bN-mAbs) to gp120 as well as studies of the carbohydrate content of virion associated Env proteins. Beginning in 2009, studies of bN-mAbs isolated from HIV infected subjects revealed that many recognized unusual glycan dependent epitopes requiring high-mannose glycans that are early intermediates in the N-linked glycosylation pathway (11-20). Passive transfer studies reviewed by Stephenson & Barouch (21) confirmed that these bN-mAbs could protect animals from infection by SHIV viruses (22-27) and lower virus loads in HIV infected individuals (28, 29). Multiple studies have now demonstrated that the carbohydrate present on virion associated envelope glycoprotein, representing approximately 50% of its molecular weight, is enriched for simple, high-mannose forms of N-linked carbohydrates rather than the complex, sialic acid containing glycans found on most membrane bound and secreted glycoproteins (20, 30-32). Since the rgp120 vaccine used in the RV144 study and other clinical trials (33-35) was enriched for complex glycans (36), they lacked multiple epitopes targeted by the high-mannose specific bN-mAbs. Thus the possibility exists that rgp120s such as A244-rgp120 used in the RV144 trial, produced with the glycans required to bind bN-mAbs, might be more effective in eliciting a protective immune response than the previous rgp120 vaccines. To test this hypothesis in human clinical trials, a practical way to produce large quantities of Env proteins possessing the high-mannose glycans is required.

The production of recombinant HIV envelope proteins (rgp120 and rgp140) for clinical research and commercial deployment has historically been challenging. Not only is it labor intensive to isolate stable cell lines producing commercially acceptable yields (e.g. >50 mg/mL) but it is also difficult to consistently manufacture a high quality, well defined product with uniform glycosylation. Replacement of the native envelope signal sequence (37, 38) and codon optimization (39) improved yields, but generating stable CHO cell lines suitable for vaccine production remained difficult. Consequently, the antigens used in the RV144 trials manufactured more than 20 years ago are still being used in multiple clinical trials (40-44).

Recombinant gp120 typically possesses 25 or more potential N-linked glycosylation sites making up more than 50% of the protein’s mass (7, 45). Each glycosylation site can possess as many as 4 sialic acid residues, with up to as 79 different glycoforms (46) possible at a single site, resulting in enormous heterogeneity in net charge and biophysical properties. This variability makes it hard to purify and define the precise chemical structure of the recombinant protein. As pharmacokinetic and pharmacodynamic properties of glycoproteins are in large part determined by the sialic acid content, glycan heterogeneity represents a major source of product variability (47).

## 2 Results

Efforts to produce HIV Env proteins for clinical testing have been complicated by problems of poor expression, heterogeneity in N-linked glycosylation and net charge, and low yields from downstream purification (36, 46, 48-56). To address these problems we combined a high efficiency electroporation device (MaxCyte STX, MaxCyte Inc., Gaithersburg, MD), a robotic cell selection system, ClonePix2 (Molecular Devices, Sunnyvale, CA) and a novel cell line (MGAT1^-^ CHO 3.4F10) that was recently developed in our lab (57). The MGAT1^-^ CHO cell line has a mutation in the Mannosyl (Alpha-1,3-)-Glycoprotein Beta-1,2-N-Acetylglucosaminyl-transferase gene (MGAT1) introduced by CRISPR/Cas9 gene editing. Recombinant gp120 produced by transient transfection in MGAT1^-^ CHO exhibited enhanced binding to three major families of glycan dependent bN-mAbs (PG9, PGT128, and PGT121/10-1074) compared to rgp120s produced in normal CHO or 293 cell lines. To explore the utility of MGAT1^-^ CHO cells as a cellular substrate for biopharmaceutical manufacturing of HIV vaccines, we attempted to create a stable MGAT1- CHO cell line expressing a variant of the A244-rgp120 envelope protein that was used in the RV144 HIV-1 vaccine trial (1). This variant, A244_N332-rgp120, differed from the A244-rgp120 immunogen in that a single N-linked glycosylation site was moved from N334 to N332 (58).

### 2.1 Replacement of chemical transfection with electroporation

Estimating that the frequency of cells expressing high levels of rgp120 might be in the range of one in 10^−4^ (59-61), we calculated that we needed to screen 10- 100 thousand transfectants. To optimize transfection efficiency, we replaced cationic lipid transfection that we had previously used to transiently produce rgp120’s (36, 57, 58, 62) with electroporation. Use of the MaxCyte STX system resulted in reproducible transfections with efficiencies typically greater than 80% in CHO-S or MGAT1^-^ CHO cells when GFP expression was quantified by flow cytometry, and viabilities greater than 95% measured by trypan-blue exclusion (Fig. 1). Based on these results, MGAT1^-^ CHO cells were electroporated with a plasmid designed for the expression of A244-N332-rgp120, and the aminoglycoside 3’-phosphotransferase gene that confers resistance to selectable marker, G418.

**Figure 1.**
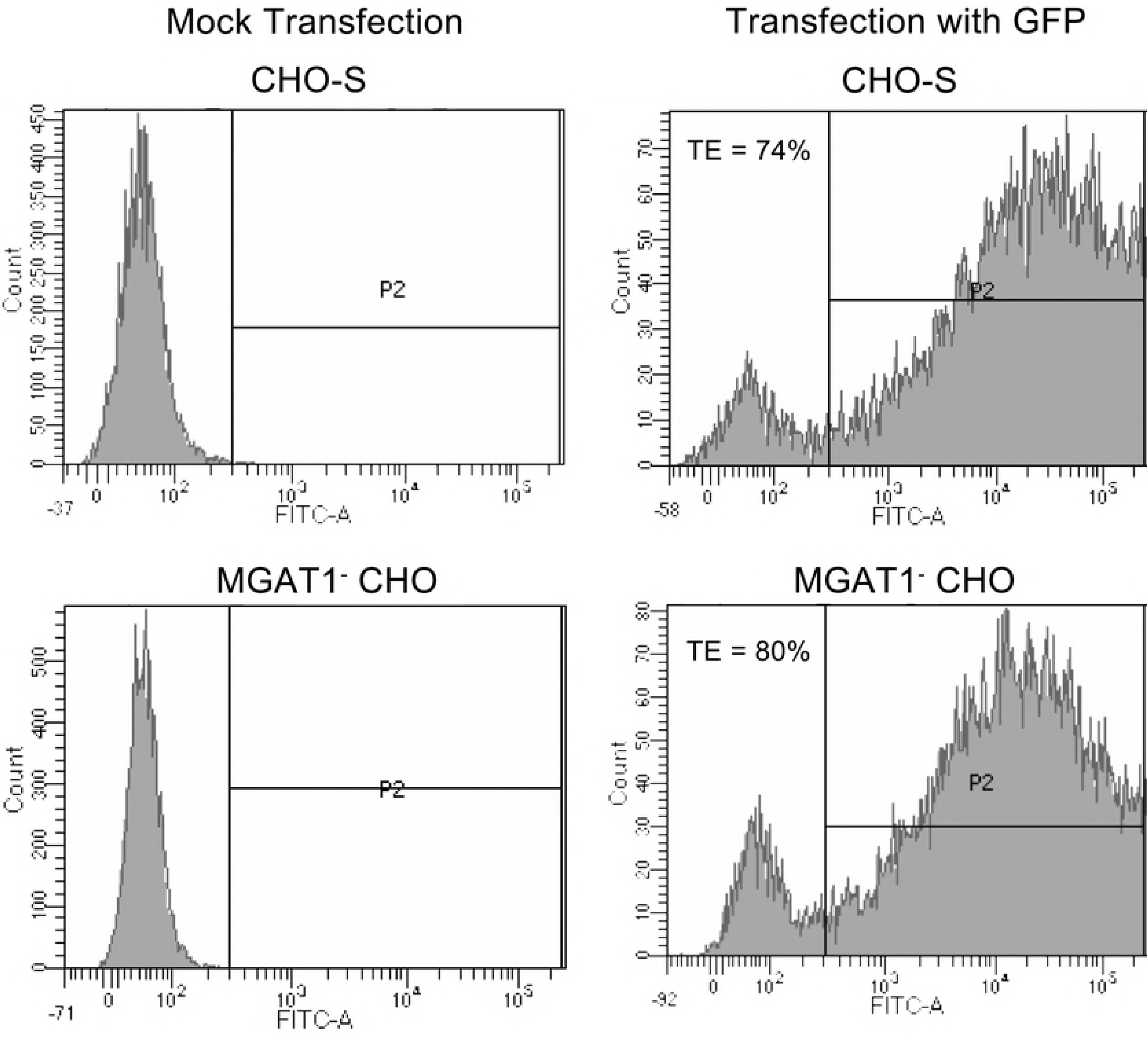
Transfection efficiency of MGAT1-CHO-S cells as determined by expression of green fluorescent protein (GFP). MGAT1^-^ CHO and CHO-S cells were transfected by electroporation with a linearized green fluorescent protein expression plasmid or mock electroporated. Forty-eight hours after transfection, viability was determined by Trypan blue exclusion on a BioRad TC10 as >95% by for both lines. Flow cytometry on a LSRII (Becton Dickinson, San Jose, CA) was used to calculate the percentage of transfected cells expressing GFP.

### 2.2 Selection of stable MGAT1^-^ CHO cell lines expressing A244_N332-rgp120 envelope proteins

We used the ClonePix2 cell screening and selection robot to identify and select the comparatively few transfectants secreting high levels of rgp120. In this system, high producing cell colonies are visualized by the formation of a “halo” or immunoprecipitin band formed by fluorescently labeled antibodies suspended in a semi-solid, methylcellulose containing, cell culture matrix. After electroporation, cells were suspended in a semisolid matrix containing the selectable marker G418 and antibodies to rgp120 labeled with the Alexa 488 fluorophore. After six days, distinct colonies were visible. By sixteen days, 45,000 colonies dispersed among 8 six-well cell culture plates had grown sufficiently for robotic screening and selection (Fig. 2A). When viewed under fluorescent light (Fig. 2B), a small fraction of the cells exhibited halos resulting from antibody-antigen precipitin bands that formed around the colonies secreting high levels of rgp120. The top 0.1% (44) of colonies selected based upon morphology and halo intensity were picked by the robot and expanded for further analysis (Fig. 2C). Selected colonies were subsequently screened by ELISA for the ability of secreted rgp120 to bind polyclonal antibodies, and the prototypic, glycan dependent bN-mAb PG9. PG9 recognizes an epitope in the V1/V2 domain and requires Man5 at position N160 for binding (18). Based on the ELISA results, cells from the top 25 rgp120 producing colonies were transferred to 24 well plates and screened by ELISA and immunoblot. The best 15 lines were then expanded into 125 mL cultures for quantitative protein expression assay under the same expression conditions used for transient expression i.e. CD-CHO opt-CHO supplemented with glucose, glutamine, CHO- Feed A and yeastolate. Six of these cultures were expanded for growth in 1.6 L shake flask cultures. Immunoblot analysis (Fig. 3A) revealed that rgp120 made by the 6 MGAT1^-^ CHO colonies were smaller in size (85 kDa) compared to A244-rgp120 produced in normal DG44 CHO cells (120 kDa). Comparison of reduced and non-reduced proteins detected a trace amount of aggregated rgp120 protein and no proteolytic degradation (clipping). Cultures were harvested when cell viability dropped to 50%. When culture supernatant was assayed by ELISA (Fig. 3B) rgp120 titers of approximately 400 mg/L were observed in in two cell lines (5D and 5F) with the 5C line exhibiting the lowest rgp120 titer (approximately 125 mg/L). Examining the kinetics of cell growth and rgp120 accumulation in cell culture medium (Fig. 3C), we found that rgp120 production increased after the addition of sodium butyrate at day six with the rate of accumulation stabilizing between 10-14 days of cell culture. During this period, rgp120 became the major protein secreted into the cell culture medium.

**Figure 2.**
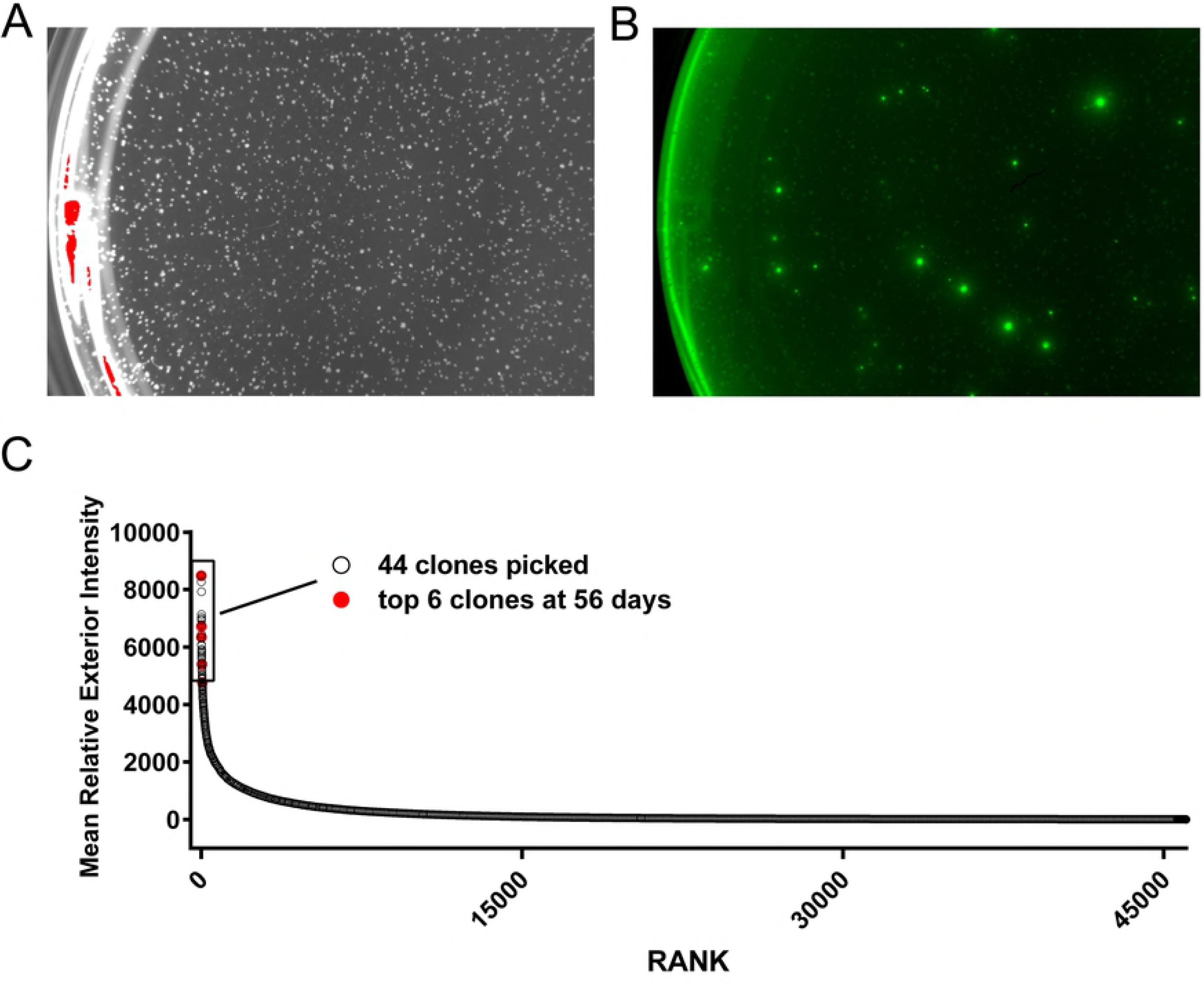
Primary identification of high producer MGAT1^-^ CHO lines expressing A244_N332 rgp120 by immunofluorescent labeling. **(A)** G418 selected colonies visible in a single 35mm well illuminated with white light at 6 days. **(B)** The same single 35mm well illuminated with 490 nm wavelength light. Colonies actively secreting rgp120 have a green “halo” visible at 525 nm. **(C)** Relative mean exterior fluorescence of halo for more than 10,000 colonies imaged by the ClonePix2 plotted by rank. The top ranking 0.1% of colonies (44) were robotically picked and cultured. The six clones expressing 0.2-0.4 g/L at day 56 are shown in red.

**Figure 3.**
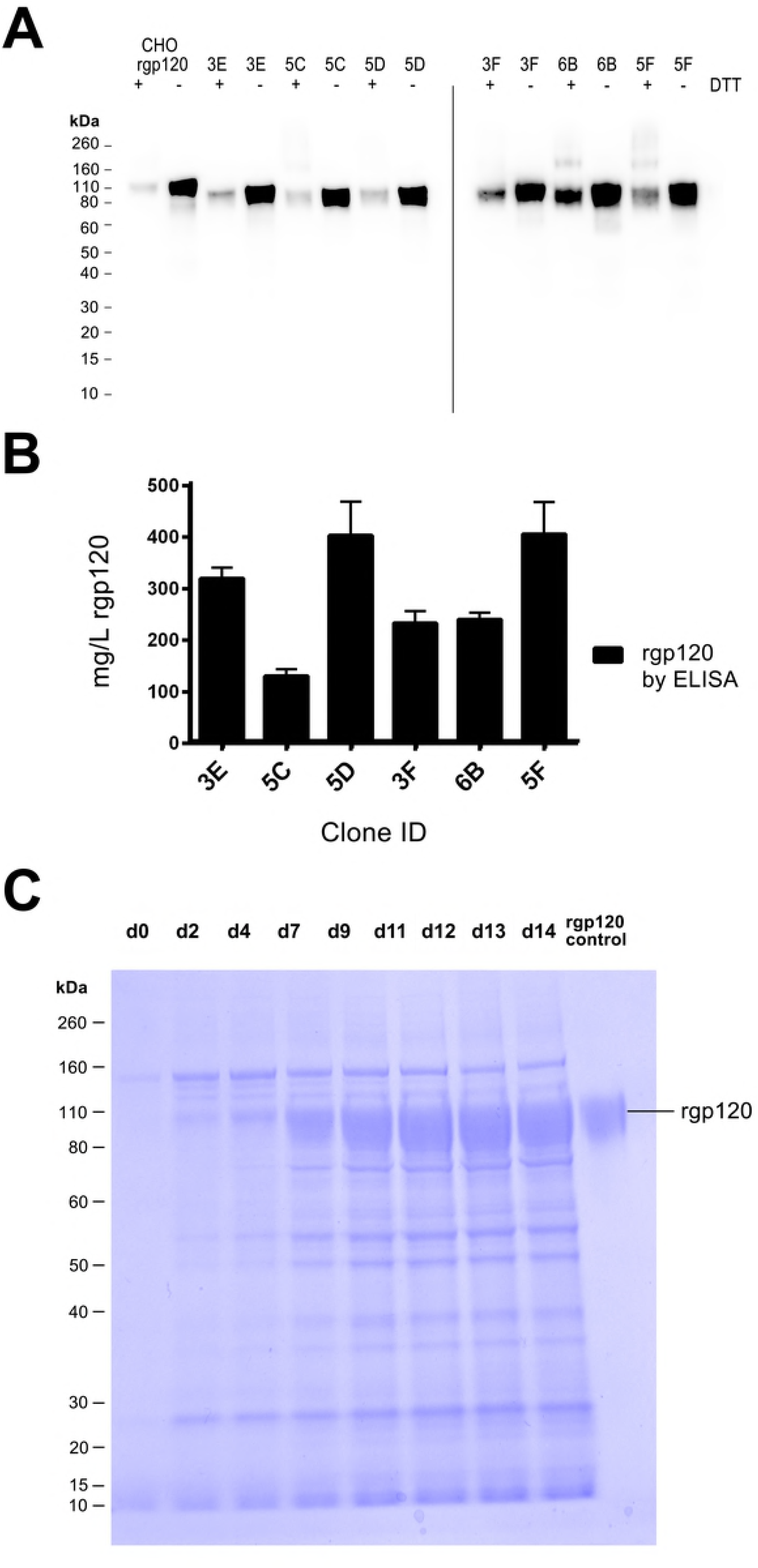
Analysis of A244_N332-rgp120 secreted from stable MGAT1^-^ CHO cell lines. Six stable MGAT1^-^ CHO cell lines identified with the ClonePix2 were selected as potential substrates for HIV vaccine production. **(A)** Immunoblot of affinity-purified rgp120 (50 ng per lane) produced by each of six A244_N332-rgp120 cell lines: 3E, 5C, 5D, 3F, 6B, and 5F. Purified A244_N332-rgp120 produced in normal CHO DG44 cells (692) was shown for purpose of comparison. **(B)** Comparison of A244_N332-rgp120 protein yields as determined by ELISA from the six MGAT1^-^ CHO cell lines. (**C)** SDS PAGE of rgp120 produced by the 5F MGAT1^-^ CHO cell line. Supernatant samples (10 μl per lane) collected over the time course of the culture were electrophoresed on a 4-12% NuPage PAGE SDS gel in MOPS buffer (Thermo Scientific, Waltham, MA). The gel was stained with Simply Blue (Thermo Scientific, Waltham, MA) and visualized using an Innotech FluoChem2 system (Genetic Technologies, Grover, MO).

### 2.3 Growth characteristics of MGAT1- CHO cells expressing A244_N332-rgp120

Several experiments were performed to characterize the growth characteristics of MGAT1^-^ CHO cells expressing A244_N332-rgp120. The initial 125-500mg/L yield, was obtained using culture conditions primarily designed for transient expression of recombinant proteins following electroporation. Data is shown in Figure 4A-C from triplicate cultures of 5F MGAT1^-^ CHO (600 mL cultures grown in 1.6 L shake flasks) for a 13-day culture period. Cells were grown at 37°C until they reached late log growth phase (six days) then sodium butyrate was added to enhance protein expression and the temperature was shifted to 32°C for the remainder of the experiment (Fig. 4A). The cell viability ranged from 90-100% for the first 8 days of culture and then steadily declined. In contrast the cumulative amount of rgp120 in the cell culture medium continued to accumulate over the entire 13-day culture (Fig. 4C) reaching a maximum of 800 mg/L by harvest. Figure 4 panels D-F show a similar batch fed experiment to test the effect of different feed additives on protein production by the 5F MGAT1^-^ CHO line. Five (duplicate) batch fed culture of the 5F MGAT1^-^ CHO isolate were grown in shake flasks in balanced CHO-Growth A (Irvine Scientific, Santa Ana, CA) media supplemented with CHO Feed C, glucose, glutamine and one of each of a panel of peptone hydrolysates; yeastolate, cottonseed, pea, wheat or CD-hydrolysate, which support protein expression in CHO cells, reviewed in (63). The cells were again grown at 37°C until they reached late log growth phase (six days) with a viable cell density approaching 1×10^7^ cells per ml, adding Sodium butyrate (1mM) and shifting the temperature 32°C for the remainder of the experiment (Fig. 4D). There were small differences in cell growth and viability (Fig. 4D and 4F) and productivity with the different peptone hydrolysate additives which might be further explored prior to large scale production, however, all supported 1g/L production of rgp120 at harvest (Fig. 4F). These studies demonstrate that the 5F MGAT1^-^ CHO cell line expressing A244_N332-rgp120 can be grown to high cell densities and is productive for up to 12-14 days in culture. It is likely that media optimization and a regulated bioreactor system can improve cell viability, cell densities, and rgp120 expression

**Figure 4.**
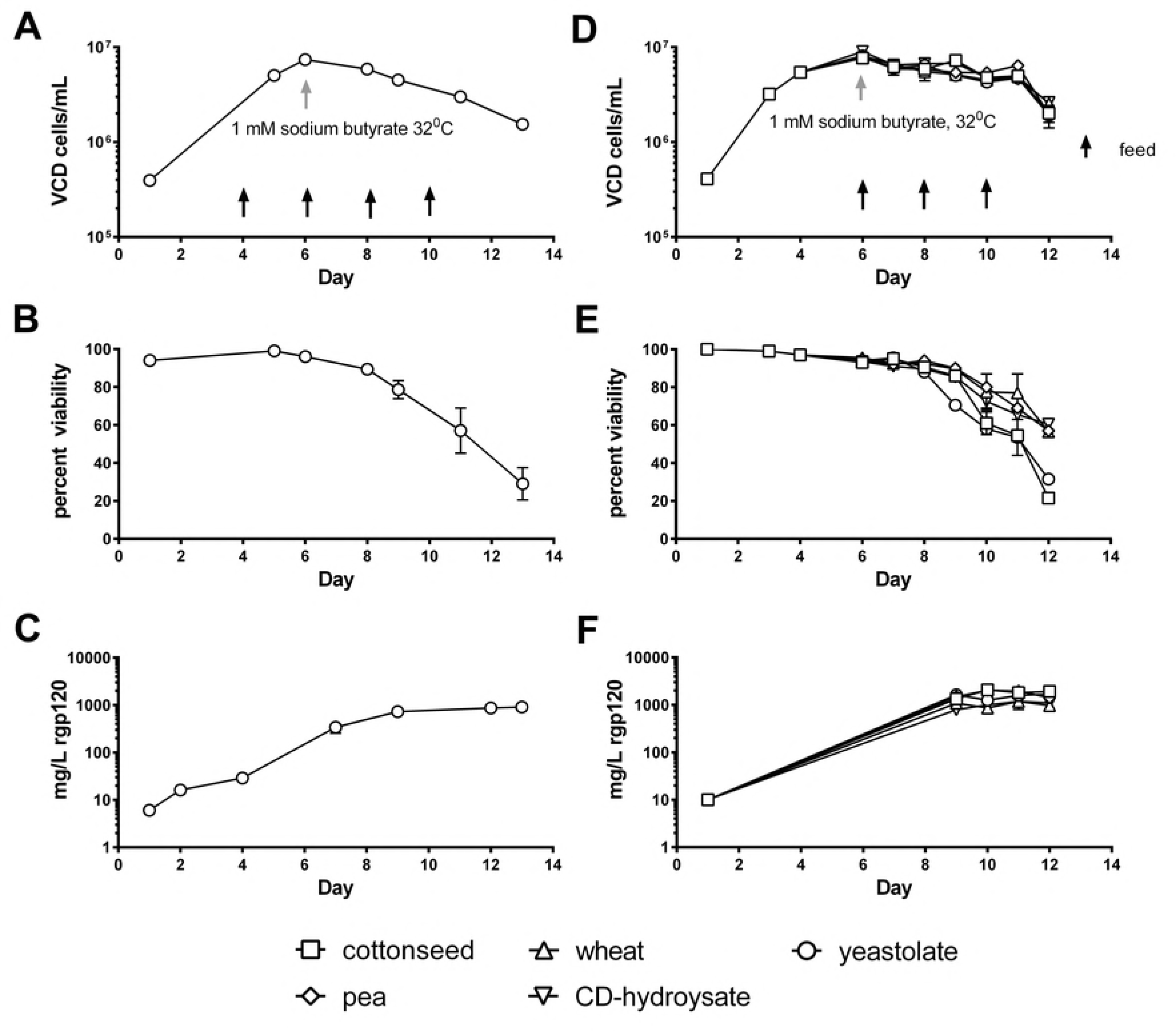
Growth and expression of the 5F MGAT1- CHO cell line expressing A244_N332-rgp120 in shake flask cultures. Cells were cultured under standard conditions until day 6 when 1 mM Sodium butyrate was added, and the temperature shifted to 32°C. Panels **A-C**: cells were fed with CHO Feed A and yeastolate as indicated, and harvested at day 13 (data from 3 shake flasks averaged). **(A)** Timecourse graph of viable cell densities (VCD) determined by trypan-blue exclusion on a BioRad T20 cell counter. **(B)** Timecourse of cell viabilities determined by trypan-blue exclusion. **(C)** Timecourse of A244_N332-rgp120 protein accumulation determined by ELISA. Panels **D-F** demonstrate optimization of protein expression (at >1g/L) by use of different feed additives. Five duplicate pairs of cultures were fed (as indicated) with CHO Feed C and either yeastolate (BD, Franklin Lakes NJ), cottonseed, wheat, pea hydrolsate (Friesland Camparia, Delhi, NY) or CD-hydrolysate (SAFC, Calsbad CA) at days 6, 8 and 10, and harvested at day 12 (data from each pair of shake flasks is averaged). **(D)** Timecourse graph of viable cell densities (VCD) determined by trypan-blue exclusion on a BioRad T20 cell counter. **(E)** Timecourse of cell viabilities determined by trypan-blue exclusion. **(F)** Timecourse of A244_N332-rgp120 protein accumulation determined by ELISA.

### 2.4 Sensitivity of A244-N322-rgp120 to digestion by Peptide-N-Glycosidase F and Endoglycosidase H

Recombinant gp120 produced in the 5F MGAT1^-^ CHO cell line exhibited an apparent molecular weight (MW) of (85 kDa). The same protein produced in CHO-S cells, exhibited an apparent molecular weight of 120 kDa (Fig. 3A). This difference in size would be expected if the glycans present in the protein produced in the MGAT1^-^ CHO cell line were limited to Mannose −5 (Man5) or earlier intermediates in the N-linked glycosylation pathway, and the glycans present in the protein produced in the CHO-S cells consisted of the normal sialic acid containing complex carbohydrates. To test this hypothesis, the proteins were digested with endoglycosidase H (EndoH) or Peptide-N-Glycosidase F (PNGase F) (Fig. 5). EndoH selectively cleaves within the chitobiose core of high-mannose and some hybrid oligosaccharides and thus cleaves the simple, high mannose forms, of N-linked glycans but not the mature sialic acid containing N-linked glycans. In contrast, PNGase F cleaves between the innermost N-acetylglucosamine and asparagine residues of high mannose, hybrid, and complex oligosaccharides and is able to digest both simple and complex N-linked glycans. We observed that PNGase F treatment converted the proteins produced in both the MGAT1^-^ CHO and CHO-S cells to a common molecular weight of approximately 56 kDa. This result confirmed that the difference in molecular weight between the proteins produced in the MGAT1^-^ CHO cell line and the CHO-S cell line could be attributed to differences in the type of glycosylation and that approximately 50% of the mass of rgp120 is carbohydrate. When the sensitivity to EndoH was measured, we found the rgp120 produced in the MGAT1^-^ cells was mostly sensitive to digestion by EndoH, whereas the rgp120 produced in the CHO-S cells was resistant to EndoH digestion. This result confirmed that rgp120 produced in MGAT1^-^ CHO cells is derivatized primarily with simple, high mannose glycans whereas the protein produced in CHO-S cells is derivatized primarily with the complex, mature form of N-linked glycosylation.

**Figure 5.**
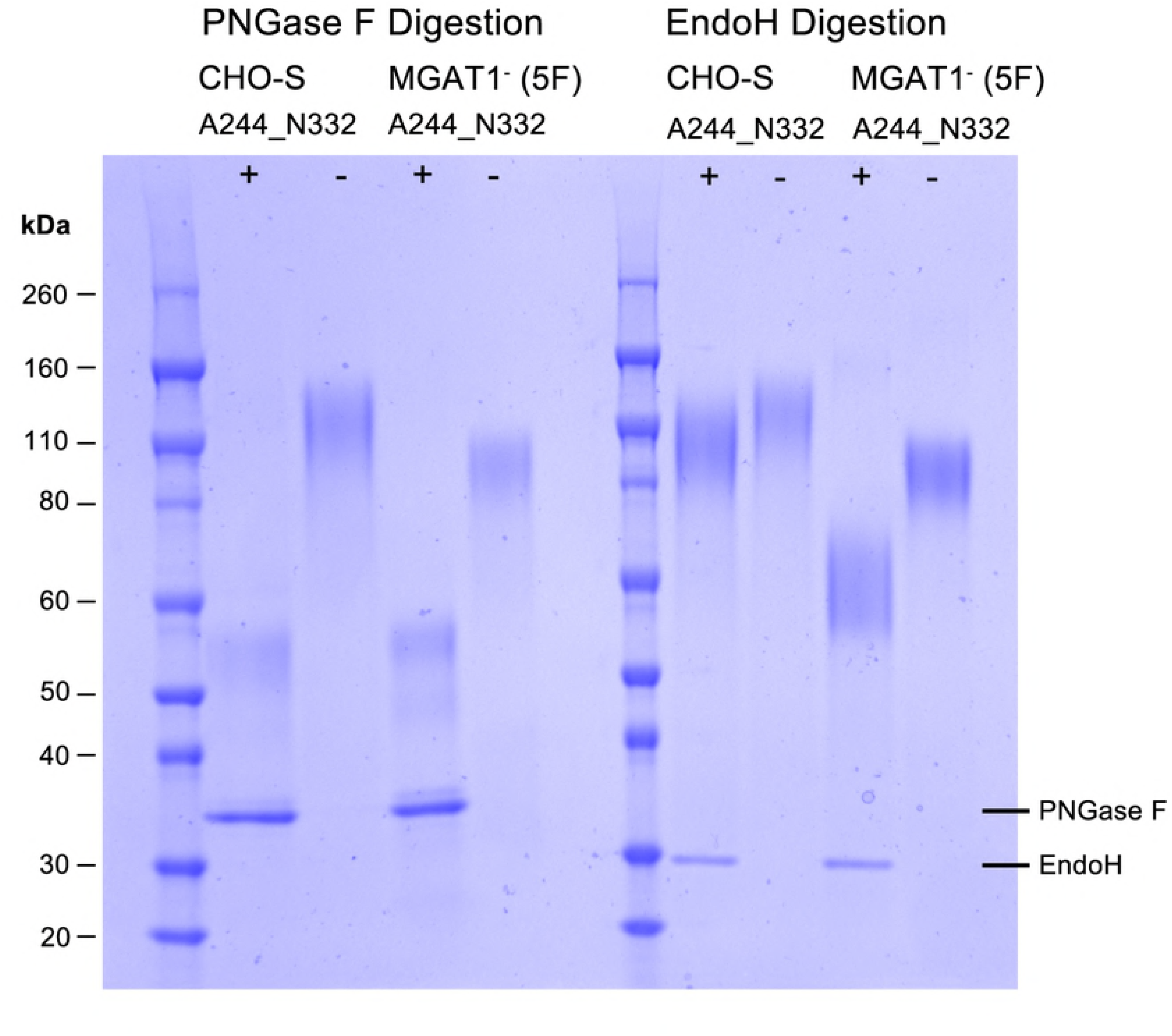
SDS-PAGE analysis of A244_N332 rgp120 HIV produced in 5F MGAT1^-^ CHO and CHO-S cells treated with PNGase or EndoH. Enzymes and buffers were purchased from (New England Biolabs, Ipswich, MA). Purified protein was denatured and reduced then incubated overnight at 37°C with or without glycosidase. Protein was resolved (2 μg/lane) on 4–12% SDS-PAGE gel and stained with Simply Blue. Plus (+) indicates enzyme treatment, minus indicates untreated.

### 2.5 Binding of A244_N332-rgp120 by bN-mAbs

The functional differences in the antigenic structures of A244_N332-rgp120 produced in the MGAT1^-^ CHO cells and A244 rgp120 produced in normal CHO-S cells was measured in a series of antibody binding experiments (Fig. 6). For these studies, the binding of bN-mAbs to rgp120s expressed in a stable MGAT1^-^ CHO cell line (5F MGAT1^-^ CHO) was compared to bN-mAb binding to the same protein expressed by transient transfection in MGAT1^-^ CHO cells, and the A244 rgp120 protein transiently expressed in CHO-S cells. In this regard, the protein expressed in CHO-S cells closely resembled the A244-rgp120 protein used in the RV144 clinical trial. The panel of bN-mAbs included both glycan dependent bN-mAbs PG9, PGT121/101074, and PGT128 (17, 18) as well as the CD4 supersite site VRC01 antibody (64, 65).

**Figure 6.**
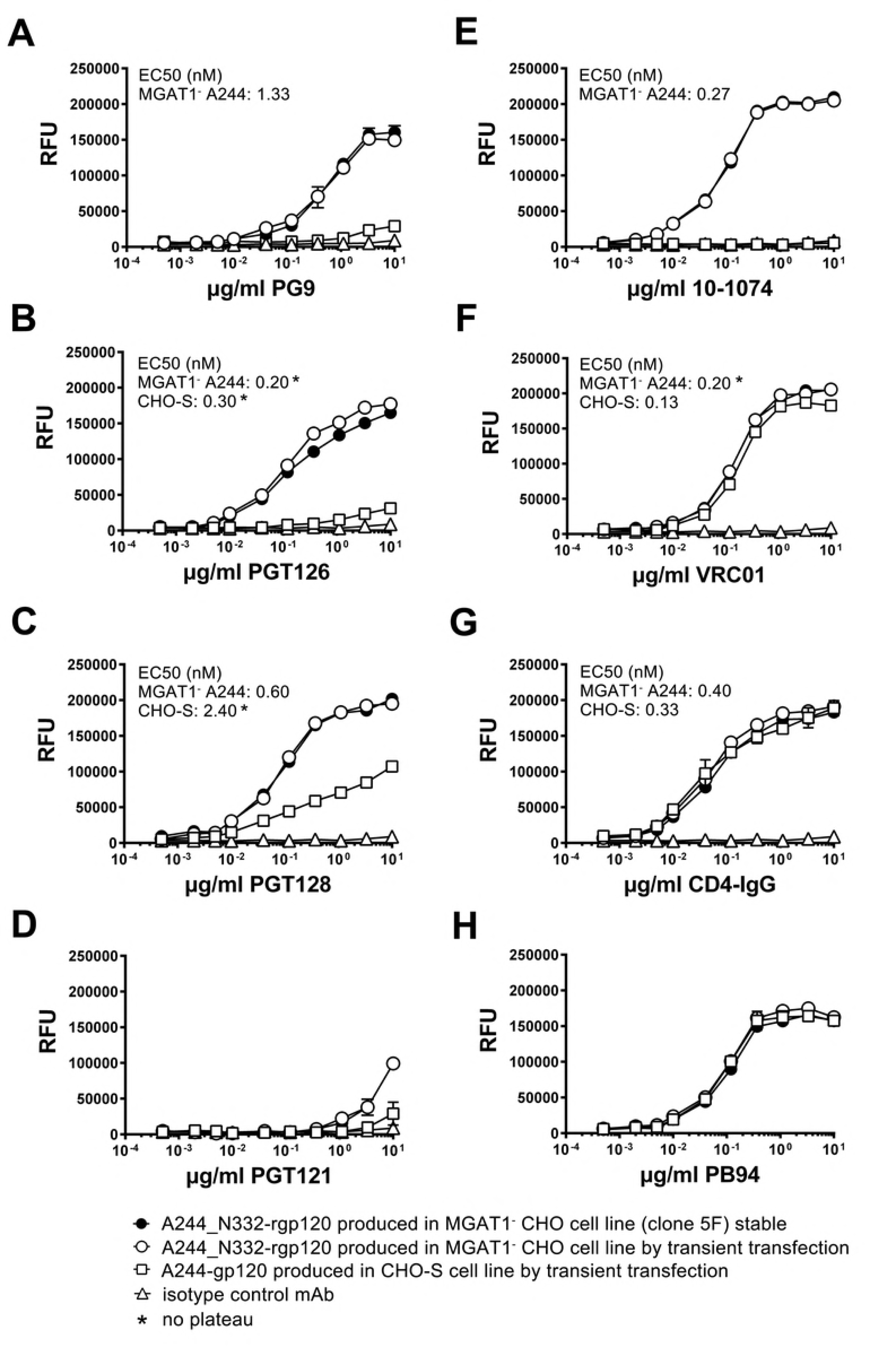
Binding of bN-mAbs to A244-rgp120 produced in normal and A244_N332-rgp120 produced in MGAT1^-^ CHO cell lines. A244_N332-rgp120 was purified from the stable clone 5F MGAT1^-^ CHO cell line (closed circles) or from the MGAT1^-^ CHO cells (open circles) transiently transfected with the UCSC 1331 plasmid. A244-rgp120 expressed and purified from transiently transfected CHO-S cells (open squares). Antibody binding was measured by a fluorescent immunoassay (FIA).

For purposes of comparison, rgp120 binding to the virus entry inhibitor CD4-IgG served as a positive control. We first examined rgp120 binding by the prototypic glycan dependent bNAb PG9, that binds the core mannose residues of two glycans at position N160 and either N156 or N173 within the V1V2 domain (18). The glycan at N160 was initially identified as Man5 in the context of a V1V2 scaffold structure (66) but recent reports suggest that PG9 is more tolerant of heterogeneity than initially thought (14, 67). Consistent with the hypothesis that a complex glycan at position N160 might interfere with PG9 binding, glycan mapping of CHO-S produced monomeric rgp120 revealed complex glycans at positions N156 and N160 (68, 69). We observe a quantitative difference (Fig. 6A) between the binding affinity of rgp120 A224-N332 produced in MGAT1^-^ CHO cells as measured by EC50 (1.33 nM) for PG9, when compared to the RV144 antigen, rgp120 A224, produced in CHO S cells (no binding plateau). This result is in concordance with data from previous transient transfection studies (57, 58). Binding of bN-mAbs to protein produced in the 5F MGAT1^-^ CHO cell line was indistinguishable to protein produced by transient transfection in MGAT1^-^ CHO cells.

We examined rgp120 binding to four bN-mAbs from two different families that are known to recognize glycan dependent epitopes in the stem of the V3 domain. PGT126 and PGT128 bN-mAbs are both members of the PGT128 family, and were derived from a common ancestral immunoglobulin VH gene (17). The bN-mAb 10-1074 is a member of the PGT121 family, and both antibodies were derived from a common ancestral immunoglobulin VH region gene distinct from the PGT128 family (13, 17). We observed significantly improved binding to members of the PGT128 family to MGAT1^-^ CHO-S produced A244_N332 rgp120 compared to the CHO-S A244 protein; however the magnitude of the difference was much greater for PGT126 compared to PGT128 (Fig. 6B and 6C). Improved PGT128 family binding is consistent with enrichment of N332 with Man5-9 glycans as PGT126/128 epitopes bind core residues of high mannose at positions N301 and N332, or N295 and N334 (15, 17, 19, 20). Mass spectroscopy of virion-derived gp120 has shown that asparagine at position 332 is occupied by Man5-9 glycans with Man8-9 dominating (32). However on CHO-S derived rgp120, high mannose is dominant but not exclusive (68, 69). Our binding data suggests that the PGT126 epitope is more sensitive to occlusion by proximal processed glycans than PGT128.

Members of the PGT121 family (PGT121 and 10-10-74) differed considerably in their ability to bind MGAT1- A244_N332 and CHO-S A244 rgp120 (Fig. 5D and 5E). PGT121 is different from all of the other glycan dependent bN-mAbs tested in that it accommodates either a sialylated or high mannose glycan at positions N332 or N334, and a sialyated glycan at position N137, whereas 10-1074 is N332 high-mannose restricted (13, 17). We observed significantly improved binding of 10-1074 to A244_N332 rgp120 produced in the MGAT1^-^ CHO cell line, and poor binding of PGT121. PGT121 did not bind rgp120 made in CHO-S cells. In control experiments, we found that all of the proteins tested bound to VRC01 bN-mAb. We noted a small but consistent higher maximal binding of VRC01 to proteins produced in the MGAT1^-^ CHO cell line compared to the CHO-S cell line (Fig. 6F). This difference was also previously reported (57). VRC01 is an anti-CD4 binding site antibody with low affinity for glycan in glycan-array assay (65). Glycans N197, N276 and N262 or N263 overlap with the binding site, and an N276 mannose core /VRC01 light chain contact was recently identified by (70). All proteins tested bound comparably to CD4-IgG regardless of the expression system (Fig. 6G), indicating that the CD4 binding site was properly folded and that the concentrations of the individual proteins were identical. Similarly, all of the Env proteins captured with the 34.1 Mab used in this experiment, bound comparably to the PB94 polyclonal rabbit sera (Fig. 6H). Thus, there was no significant difference in the concentration or coating efficiencies of the proteins used for the binding studies.

### 2.6 Pathogen testing

In order for the 5F MGAT1^-^ CHO cell line to be considered as a substrate for vaccine production by current Good Manufacturing Processes (cGMP), it needs to be free of contamination by other cell lines and adventitious agents. To obtain data supporting these criteria, cells from the 5F MGAT1^-^ CHO cell line were screened for contamination by a commercial testing laboratory (IDEXX, Inc., Columbia, MO). This analysis used validated PCR based techniques to screen for contamination by cells from multiple other species (human, mouse) and for contamination by a large number of human and animal pathogens including mycoplasma and minute virus of mice (MVM). No cellular, viral, or microbial contamination of the original research cell bank was detected (Supplemental Tables 1 and 2).

### 2.7 Outline of new process for the development of a stable MGAT1- CHO cell line expressing HIV-1 rgp120

Based on our results we were able to devise a new standardized cell line production strategy for creating stable CHO cell lines expressing rgp120 and other Env proteins in a timeline of 8-10 weeks. An outline for this new cell line development process including the experimental timeline for electroporation, colony selection, and protein expression/ purification is shown in Figure 7. The use of high efficiency electroporation, robotic screening, and the elimination of gene amplification strategies all contribute to a major reduction in the time and cost of producing stable CHO cell lines.

**Figure 7.**
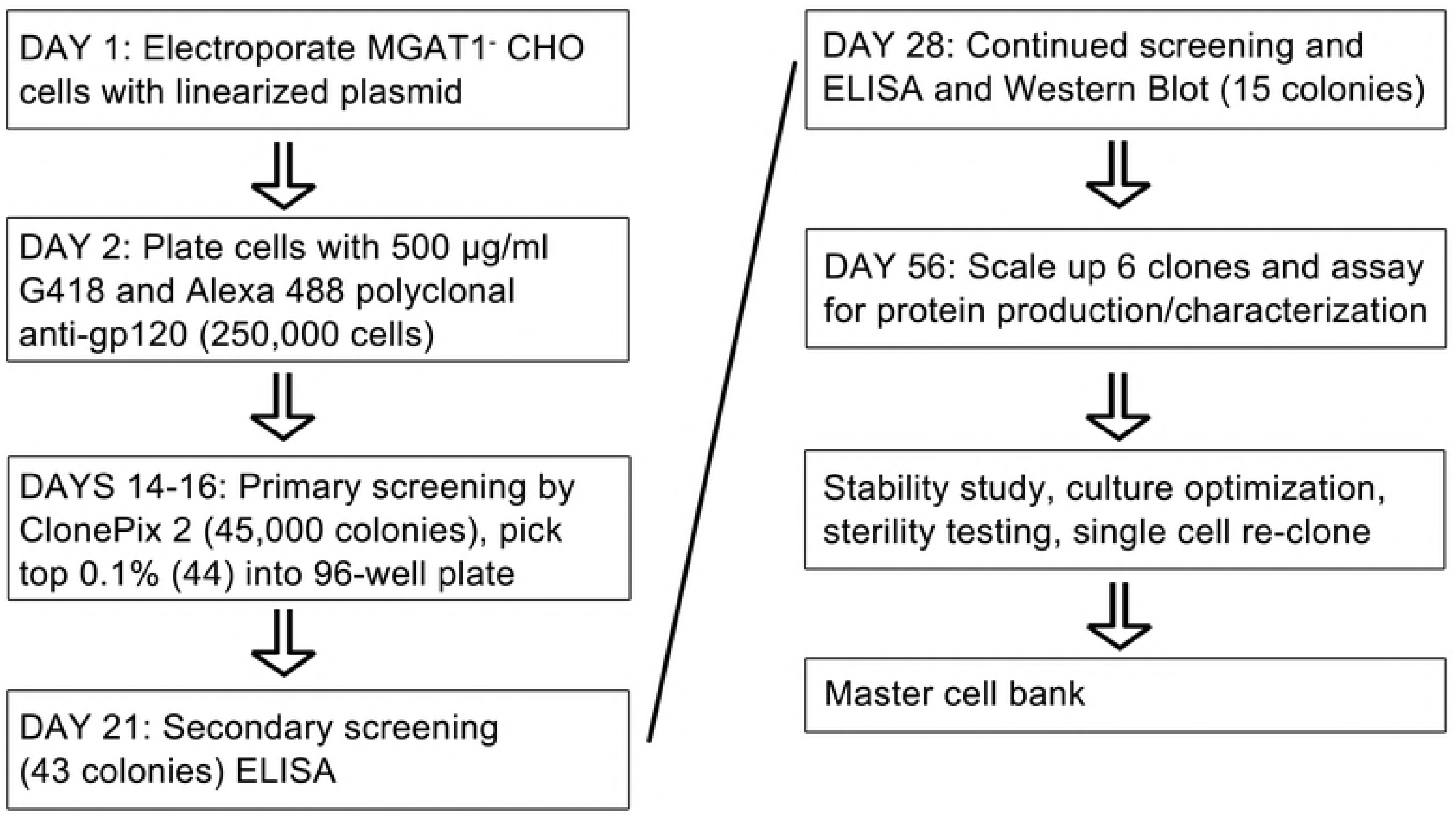
The timeline for development of stable MGAT1^-^ CHO cell lines expressing HIV-1 rgp120. Leading clones expressing 0.2-0.4 g/L in shake flasks under standard laboratory conditions were selected in less than two months. Production was subsequently increased to levels of g/L rgp120 production with minimal feed optimization.

## 3 Discussion

In this study we describe the development of a new robotic cell line screening and selection strategy for the rapid production of stable, high yielding CHO cell lines suitable for the production of HIV vaccines. We also demonstrate that this strategy can be successfully employed using the recently described MGAT1^-^ CHO cell line (MGAT1^-^ CHO 3.4F10) that limits glycosylation primarily to Man5 glycans and, to date, has only been used for transient transfection experiments (57). Finally, we describe a MGAT1^-^ CHO cell line (5F MGAT1^-^ CHO) that produces high levels of a variant of gp120 from the A244 isolate of HIV-1. A244-rgp120 was a key component in the AIDSVAX B/E vaccine (5, 6) used in the RV144 clinical trial. The Env protein (A244_N332-rgp120) produced by the 5F MGAT1- CHO cell line differs from the A244-rgp120 used in the RV144 trial in the location of a single N-linked glycosylation site (N332 compared to N334) and that it incorporates primarily Man5 glycans. These changes enable the A244_N332 Env protein to bind bN-mAbs from three major families of bN-mAbs (PG9, PGT128, and PGT121/10-1074) that did not bind to the original vaccine immunogen. The addition of these glycan dependent epitopes represents a significant improvement in the antigenic structure of A244-rgp120 and might improve the level of efficacy that can be obtained from an RV144-like immunization regimen.

Prior to the development of the MGAT1^-^ CHO 3.4F10 cell line there was no practical means by which recombinant Env proteins enriched for high mannose glycans could be produced at large scale for HIV vaccine production. In these studies, we demonstrated that CHO cell lines expressing up to 1g/L of rgp120 that are potential candidates for vaccine antigen production, can be produced within 2-3 months. In contrast, the CHO cell line used to produce A244-rgp120 used in the RV144 trial produced only 20 mg/L and took more than 18 months to develop (Berman, P.W., personal communication).

Previous studies have shown that early intermediates in the N-linked glycosylation pathway (Man5 or Man9) are essential components of many epitopes recognized by bN-mAbs (11-16, 18, 19, 20). Additionally, we now know that HIV virion-associated Env proteins are enriched for these early intermediates in the glycosylation pathway (20, 31, 50, 71). The A244_N332-rgp120 protein produced in the 5F MGAT1^-^ CHO cell line possesses multiple glycan dependent epitopes recognized by bN-mAbs and appears to possess a glycan structure that more closely resembles authentic HIV Env proteins compared to Env proteins produced in normal CHO cells. We hypothesize that this “glycan optimized” immunogen produced in the 5F MGAT1^-^ CHO cell line might increase the potential level of protection documented in the RV144 trial from 31.2% (P=0.04) to the efficacy level of 50% or more thought to be required for regulatory approval and clinical deployment (72).

These studies demonstrate that the MGAT1^-^ CHO cell line is a suitable substrate for stable cell line development (57). The 5F MGAT1^-^ CHO line can be grown at high densities (2×10^7^cells/mL) in serum free media for the length of time (8-10 days) consistent with modern manufacturing methods intended for the production of HIV vaccine immunogens. The success in cell line isolation was greatly enhanced by the use of robotic selection. The MaxCyte STX electroporation system achieved transfection efficiencies of greater than 80% with linearized plasmid. Another key step in the robotic screening strategy was the need to develop an imaging reagent that formed “halos” around rgp120 transfected cells. We found that the best “halos” resulting from the formation of immune-precipitin bands in semi-solid methylcellulose media were only obtained with fluorescently labeled polyclonal antibodies. Although previous anecdotal reports suggested that mixtures of fluorescently labeled monoclonal antibodies could form the immune-precipitin bands required for robotic selection, we were unable to confirm these reports. Finally, we attempted to see if the robotic selection could overcome the need for time-consuming gene amplification experiments (via methotrexate or methionine sulphoximine), and we found that directly screening approximately 40,000 clones in a single ClonePix2 experiment was adequate to isolate a cell line that produced approximately 1g/L of rgp120 with no gene amplification. Studies are in progress to determine whether the high expression levels found in this study results from the selection of cell lines with high levels of amplified genes or from integration of the HIV transgene into transcriptionally active regions of chromatin.

The new technology outlined here should allow for the rapid production and testing of multiple new Env based vaccine concepts that have not previously been tested for lack of a fast and cost effective manufacturing process. Such concepts include multivalent rgp120 vaccines (73), guided immunization vaccine strategies (73-77), and Env proteins optimized for the binding of inferred ancestral forms of bN-mAbs (78-80). In summary, by combining recent developments in transfection technology, robotic selection, and gene editing, we have developed a novel method for the production of recombinant Env proteins has the potential to improve the potency, shorten the time, and lower the cost of HIV vaccine production. These improvements provide the means to break the bottleneck in HIV vaccine manufacturing that has limited the field for the last 20 years (81).

## 4 Materials and methods

### 4.1 Cells and antibodies

The suspension adapted, stable MGAT1^-^ CHO cell line was created by targeted inactivation of the gene encoding the enzyme, Mannosyl (Alpha-1,3-)-Glycoprotein Beta-1,2-N-Acetylglucosaminyl-transferase in CHO cells using CRISPR/Cas9 gene editing (57). Suspension adapted CHO-S cells were obtained from Thermo Fisher (Thermo Fisher, Life Technologies, Carlsbad, CA). GnTI^-^ 293 HEK cells were obtained from the American Tissue Type Collection (ATCC). Broadly neutralizing monoclonal antibodies (bN-mAbs), PG9, PGT121, PGT126, PGT128, VRC01, and 10-1074, were obtained from the NIH AIDS Reagent Program (Germantown, MD) or produced from published sequence data. The entry inhibitor CD4-IgG was identical to that described by Capon et al. (82). The 34.1 murine monoclonal antibody was developed in our laboratory (62) and is specific for a 27 amino acid sequence of Herpes Simplex Virus Type 1 glycoprotein D (gD) that has been used previously as a purification tag (7, 38). Polyclonal antibodies were raised according to the guidelines of the Animal Welfare Act. The immunization protocol was reviewed and approved by the Animal Care and Use Committee of the Pocono Rabbit Farm and Laboratory (Pocono Laboratories and Rabbit Farm, Canadensis, PA). Polyclonal rabbit-serum (PB94) was obtained from rabbits immunized with a mixture of A244 and MN rgp120 as previously described (6). Polyclonal goat-serum was raised from goats immunized with a cocktail of purified rgp120s from three clades of HIV (CRF01_AE, B, and C) produced in GnTI^-^ 293 HEK cells. Rabbit and goat polyclonal anti-gp120 for use in immunoassays were purified by affinity chromatography using a HiTrap Protein G column (GE Healthcare, Little Chalfont, United Kingdom). Immunoaffinity purified anti-gp120 for use in the ClonePix2 robot (Molecular Devices, Sunnyvale, CA) was isolated from goat sera by immunoaffinity chromatography using a column consisting of gp120 coupled to Sepharose 4B (GE Healthcare, Little Chalfont, United Kingdom). Immunoaffinity purification involved successive passage over rgp120 bound affinity columns which were then washed with 10 column volumes of 50 mM Tris, 0.5 M NaCl, 0.1 M TMAC (tetramethyl-ammonium chloride) buffer (pH 7.4). Bound antibody was eluted with 0.1 M sodium acetate buffer, pH 3.0, and the eluent neutralized by the addition of 1.0 M Tris (1:10 ratio). The purified antibody was adjusted to a final concentration of 1-2 mg/mL in PBS buffer as determined by bicinchoninic acid (BCA) and conjugated to Alexa 488 (Thermo Fisher Scientific, Waltham, MA), before 0.1 μM filtration.

### 4.2 Cell culture conditions

MGAT1^-^ and CHO-S cells were maintained in CD-CHO medium supplemented with 8 mM GlutaMAX, 0.1 mM Hypoxanthine, and 0.16 mM thymidine (HT) in shake flasks using a Kuhner ISF1-X shaker incubator (Kuhner, Birsfelden, Switzerland) at 37°C, 8% CO_2_, and 125 rpm. Static cultures were maintained in 6, 24, and 96 well cell culture dishes (Greiner, Kremsmünster, Austria) and grown in a Sanyo incubator (Sanyo, Moriguchi, Osaka, Japan) at 37°C and 8% CO_2_. For protein production, CD-OPTI-CHO or CHO Balanced Growth A medium (Irvine, Santa Ana, CA) was supplemented with 2 mM GlutaMAX, HT and MaxCyte CHO A Feed which is comprised of 0.5% Yeastolate, BD, Franklin Lakes, NJ; 2.5% CHO-CD Efficient Feed A, 2 g/L Glucose (Sigma-Aldrich, St. Louis, MO) and 0.25 mM GlutaMAX). Cell culture media and additives were obtained from Thermo Fisher Life Technologies (Carlsbad, CA) unless otherwise stated. In preliminary batch fed cell culture experiments to optimize protein yield, we tested out an additional four peptone hydrolysates replacing yeastolate with; Proyield Cotton, Proyield Pea, Proyield Wheat (Friesland Campira, Delhi, NY) and CD-Hydrolysate (SAFC, Carlsbad, CA) and CHO-CD Feed efficient A with CHO-CD Efficient Feed C (Thermo Fisher Life Technologies, Carlsbad, CA).

### 4.3 Molecular cloning and sequencing

Standard genetic engineering techniques were used to construct a G418 selectable expression vector (UCSC1331) that encodes gp120 from the clade CRF01_AE strain of HIV-1. The protein produced is identical in sequence to the A244-rgp120 protein used in the RV144 clinical trial with the exception that a single N-linked glycosylation site at N334 has been moved to position N332, described by Doran et al. (58), GenBank ref MG189369. The plasmid was similar to the commercially available pCDNA3.1 vector with the exception that methylation targets at positions C41 and C179 in the CMV promoter were deleted as described by Moritz and Gopfert (83). All sequencing was performed at the University of California Core Sequencing Facility (Berkeley, CA). pCI_GFP, a gift from Dr. James Brady (MaxCyte), was transfected in parallel with the gp120 expression plasmid to monitor transfection efficiency. Plasmid DNA was prepared using the endotoxin free Qiagen Giga Prep purification kit (Qiagen, Hilden, Germany) and linearized by digestion with Pvu1 (New England Biolabs, Ipswich, MA) prior to electroporation.

### 4.4 Selection of stable MGAT1^-^ CHO cell lines expressing A244-rgp120

Electroporation of the UCSC1331 plasmid into MGAT1- CHO cells was performed using a MaxCyte scalable transfection system (STX, MaxCyte Inc., Gaithersburg, MD) according to the manufacturer’s instructions. Twenty-four hours after electroporation, cells were diluted to a concentration of 1000 or 5000 cells/mL in methylcellulose CHO-Growth A with L-glutamine (Molecular Devices, Sunnyvale, CA) containing 500 μg/mL of G418 and 10 μg/mL of Alexa 488 labeled immunoaffinity purified polyclonal goat anti-gp120 antibody. The plates were incubated at 37°C with 8% CO_2_ and 85% humidity for 16 days, then colony selection was performed using a ClonePix2 robot (Molecular Devices, Sunnyvale, CA). Colonies were imaged under white and fluorescent light (470 nm excitation and 535 nm emission wavelength filter set). Both images were superimposed, and colonies selected according to mean exterior fluorescent intensity (84). The top ranking 0.1% of colonies were aspirated with micro-pins controlled by the ClonePix2 system and dispersed automatically in a 96-well plate containing CHO Growth A medium (Genetix Molecular Devices, Sunnyvale, CA) supplemented with HT, 8 mM GlutaMAX, and 500 μg/ml G418, and cultured at 37°C, with 8% C0_2_ and 85% humidity. After 9 days in culture, protein production was assayed by ELISA, and positive colonies transferred to 2 mL wells and shake flasks when cell mass permitted transfer. Six lines were cultured for protein production (Section 2.2) at a volume of 600 mL in 1.6 L shake flasks (Thompson, Oceanside, CA)

### 4.5 Protein quantification

Protein concentration was determined by capture ELISA (62). Purified protein and cell culture supernatant were analyzed on 4-12% Bis-Tris PAGE SDS gels in either MES or MOPS gel running buffer (Thermo Scientific, Waltham, MA). For Immunoblot, proteins were electrophoresed on a 4-12% NuPage PAGE SDS gel in MES running-buffer, transferred to a PDVF membrane, then probed with a polyclonal rabbit anti-rgp120 antibody (PB94) and an affinity purified secondary HRP conjugated goat anti-rabbit H+L chain antibody (Jackson ImmunoResearch, West Grove, PA) and visualized using an Innotech FluoChem2 system (Genetic Technologies Grover, MO).

### 4.6 Affinity purification of A244_N332-rgp120

Recombinant proteins were immunoaffinity purified from cell culture media using the gD purification tag as previously described (36) and protein concentrations determined using bicinchoninic assay (BCA).

### 4.7 Binding of bN-mAbs

The binding to bN-mAbs to purified rgp120s from the MGAT1^-^ CHO and CHO-S lines was assayed with a capture Fluorescence Immunoassay (FIA) as described previously (58). Briefly, Fluotrac high binding 96 well plates (Griener Bio-One Kremsmünster, Austria) were coated with 2ug/ml 34.1 Mab overnight in PBS, then blocked with 1%BSA/PBS 0.05% tween for 2 hours. Purified rgp120 was captured at 6ug/ml overnight at 4°C. Three-fold serial dilutions of antibody, entry inhibitor or isotype control were added to each well followed by Alexa 488 labelled polyclonal anti-species antibody (Jackson ImmunoResearch, West Grove PA). Incubations were performed for 90 min at room temperature followed by a 4x wash in PBS 0.05% tween buffer unless otherwise noted. Absorbance was read using an EnVision Multilabel Plate Reader (PerkinElmer, Inc Waltham, MA) using a FITC 353 emission filter and FITC 485 excitation filter. Assays were performed in triplicate. EC50 was calculated from a plot of log (agonist) vs response –variable slope (four parameters) on Graph Pad Prism 6 for Mac., GraphPad Software, La Jolla, CA.

Binding assays were carried out in triplicate.

### 4.8 Enzymatic digestion of carbohydrate

Enzymatic digestion of rgp120 was performed as described by Yu et al (36). For molecular mass analysis post digestion, 2 μg of each protein was electrophoresed on a 4-12% Bis-Tris PAGE SDS gel in MOPS running buffer, and stained with Coomassie Simply Blue (Thermo Scientific, Waltham, MA).

## Acknowledgements

This research was supported by NIH grant RO1AI113893. We wish to thank Chelsea Didinger for expert assistance in the preparation of this manuscript. We also wish to thank Bari Holm Nazari for assistance with flow cytometry.

